# Anesthesia is a Potent Determinant of Ultra-High Dose Rate Sparing in the Murine Total Abdominal Irradiation Model

**DOI:** 10.1101/2025.07.15.664940

**Authors:** Armin D. Tavakkoli, William W. Daley, David I Hunter, Beverly A. Allen, Gretchen C. Carpenter, David J. Gladstone, Brian W. Pogue, P. Jack Hoopes

## Abstract

Radiation therapy is a mainstay of treatment for numerous gastrointestinal (GI) malignancies, where our ability to deliver dose to tumors is limited by acute GI toxicity. Ultra-high dose-rate (UHDR) ‘FLASH’ irradiation can spare normal tissue, yet its dependence on physiological variables remains incompletely defined. We compared FLASH and conventional dose-rate (CDR) 9 MeV electron total abdominal irradiation (TAI) in C57BL/6 mice anesthetized with either intraperitoneal ketamine/xylazine or inhaled isoflurane in room air, deliberately omitting supplemental oxygen. Single doses of 14 or 16 Gy were delivered, and normal-tissue injury was quantified by time-to-25% body-weight loss.

At 14 Gy, UHDR under K/X produced a marked survival advantage: by day 14, 80% of animals had not reached the weight-loss endpoint versus 40% after CDR K/X; no FLASH benefit was discernible with ISO anesthesia. Raising the dose to 16 Gy accentuated these trends; 40% of UHDR K/X mice were still below the endpoint at study termination, whereas all CDR K/X mice met it by day 7. Again, ISO abolished sparing at both dose rates. To probe mechanism, intraperitoneal oxygen tension was measured with an optical reporter in six mice. ISO anesthesia yielded significantly higher pO_2_ (62 ± 4 mmHg) than K/X (26 ± 10 mmHg), a 2.5-fold difference.

These findings identify anesthetic-dependent oxygenation as a reproducible confounder in pre-clinical FLASH studies: elevated pO_2_ under ISO negates abdominal sparing, whereas K/X preserves it across two clinically relevant doses. Rigorous control and reporting of factors that alter tissue oxygenation are therefore essential when designing experiments and, ultimately, translating FLASH radiotherapy.

## Introduction

Radiation therapy (RT) is employed in roughly one-half of all cancer cases and contributes to about 40% of curative treatments worldwide^1^. The recent arrival of ultra-high-dose-rate (UHDR, “FLASH”) RT has reinvigorated efforts to increase the therapeutic ratio by preferentially sparing normal tissue while maintaining tumor control ^2^. Elegant proof-of-principle experiments in skin, lung and brain have demonstrated dramatic reductions in toxicity when doses are delivered in micro-second pulses at ≥ 30 Gy s^−1^ compared with conventional dose rates (CDR) ^3^. Yet extension of these findings to the abdomen—a site where acute gastrointestinal (GI) syndrome remains a foremost dose-limiting toxicity—has been inconsistent. Whole-abdomen electron FLASH protects crypt stem cells and improves survival in some reports, whereas pencil-beam-scanned proton FLASH and synchrotron proton beams have produced neutral or even worse outcomes relative to CDR irradiation ^4,5^. A potential explanation is that the FLASH effect is particularly sensitive to biological context and experimental nuance that is not being controlled for across studies.

A unifying hypothesis for FLASH sparing invokes transient oxygen depletion: extremely intense pulses have been postulated to consume dissolved O_2_ faster than it can be replenished, driving tissue into a radioprotective hypoxic state for the brief period in which radical-mediated damage is fixed ^6–9^. Bulk tissue measurements have refuted this idea through systematic study of skin, and tumor, however still debate exists on whether there could be regional or micro-localized hypoxia induced during the irradiation in FLASH. However also the data seems to be dependent upon the total dose as well as the tissue oxygen, and so there seems to be both a minimum and a maximum dose threshold to see the FLASH effect ^10,11^. It seems likely that oxygen may be a factor in this complex range where FLASH tissue sparing can be seen. If it is true that oxygenation level affects the observation, then any variable that alters baseline or dynamic tissue oxygenation could modulate—or even mask—the FLASH effect. Two variables stand out in small-animal work: the choice of anesthetic and the composition of the carrier or supplemental gas.

Intraperitoneal ketamine/xylazine (K/X) and inhaled isoflurane dominate murine RT studies because they are facile and inexpensive. Unfortunately, they create very different physiological milieus. K/X produces profound bradycardia, respiratory depression, and peripheral vasoconstriction, leading to arterial hypoxemia and tissue pO_2_ values as low as 15 mm Hg in skin and brain. Isoflurane, in contrast, raises heart rate, increases tidal volume, and vasodilates resistance vessels; when delivered in room air it maintains skin pO_2_ near 25–30 mm Hg, and when delivered in 100 % oxygen it can drive tissue pO_2_ above 50 mm Hg. Carrier-gas oxygenation matters independently of the volatile agent: supplemental oxygenation during irradiation sharply increases murine skin P_O2_ values and negates the FLASH effect ^12–14^.

Despite these well-documented effects, anesthetic protocols are rarely reported in pre-clinical FLASH papers, and when they are, oxygen supplementation is common. In a recent systematic review of electron FLASH in vivo studies, fewer than one-third specified the fraction of inspired oxygen, and none quantified tissue pO_2_. Recent results showed that this omission is not benign: in murine hind-leg skin, breathing 100 % oxygen during isoflurane anesthesia abolished FLASH sparing, whereas room-air anesthesia preserved it; moreover, female mice—whose dermis exhibited higher pO_2_ than males—ulcerated sooner after UHDR irradiation ^12,15^. These data strengthened the link between oxygen tension and FLASH, but they also raised a critical question for abdominal irradiation: does the anesthetic itself, independent of inspired O_2_, dictate whether gut tissues experience protection or injury at UHDR?

The small intestine is among the most radiosensitive organs, and weight-loss kinetics as well as crypt-regeneration assays are gold-standard read-outs for acute GI toxicity. Several groups have attempted to exploit FLASH in this setting, yet outcomes have ranged from substantial sparing to overt harm. None of these studies compared different anesthetic regimens head-to-head or measured intraperitoneal oxygenation. Given that intestinal perfusion is highly responsive to vasoactive and respiratory changes, and that K/X and isoflurane exert opposite effects on both, anesthetic selection may be a hidden confounder driving inter-study variability.

Here we directly address this gap by evaluating FLASH sparing in the mouse total abdominal irradiation model, anesthetized with either KX or isoflurane in room air. To mechanistically link any observed differences to oxygen availability, a parallel cohort received intravenous PdG4 Oxyphor and real-time phosphorescence lifetime imaging (OxyLED) was used to record intra-abdominal pO_2_ immediately after laparotomy ^16^.

## Methods

### Animals & Irradiation

All animal experiments were approved by the Dartmouth College Institutional Animal Care and Use Committee (IACUC). Eighty (N=80) C57BL/6 mice were ordered from Jackson Laboratories and allowed to acclimate to the vivarium for at least 2 weeks. Following the acclimation period, mice were weighed for 3 days to establish baseline weight. On the day of irradiation, mice were anesthetized using either an IP injection of Ketamine/Xylazine (ketamine 100 mg/kg, xylazine 10 mg/kg) or inhaled isoflurane anesthesia delivered in room air through a non-rebreather mask (3% induction for 3 minutes, then maintained on 1.5%; n=10). A Mobetron linear accelerator (LINAC; IntraOp, Inc, USA) was used to deliver 14 or 16 Gy of 9 MeV electron CDR and UHDR radiation to a 3 cm x 4 cm area centered on the mouse abdomen with a 1 cm air gap. CDR irradiation was delivered at a dose rate of 0.1 Gy/s, while UHDR was delivered at an average dose rate of 420 Gy/s. For the 14 Gy UHDR treatment, the source-to-surface distance (SSD) was set to 37 cm, and each pulse had a width of 3.10 µs delivered at a pulse repetition frequency (PRF) of 90 Hz. A total of four pulses were administered, with each pulse depositing 3.5 Gy, resulting in the prescribed 14 Gy dose. For the 16 Gy UHDR treatment, the SSD was set to 37 cm, and each pulse had a width of 3.54 µs delivered at a PRF of 90 Hz. A total of four pulses were administered, with each pulse depositing 4 Gy, resulting in the prescribed 16 Gy dose. CDR treatments were delivered using a PRF of 30 HZ, pulse width of 1.2us, and the integrated ion chambers were utilized to deliver the prescribed dose.

### Dosimetry Verification

Prior to the day of live animal irradiation, on “verification day”, an in-house beta treatment planning program was used to estimate the necessary treatment parameters for 14 and 16 Gy dose delivery. Briefly, the planning software utilizes cutout specific output data from water-tank measurements and daily output data from LINAC quality assurance procedure to calculate an estimated pulse width, repetition rate, and number of pulses needed for a given dose. These parameters were then used to deliver both CDR and UHDR irradiation to (1) radiochromic film on solid water (EBT-XD, Ashland Inc, USA) (2) pre-calibrated FlashDiamond (uD) detector (PTW Inc., USA) placed at a depth of 1 mm in solid water, and (3) radiochromic film on the abdomen of a mouse phantom. The dose delivered was within 3% of the expected dose. The cutout depth-dose profile is presented in Figure 1.

**Figure 1:**
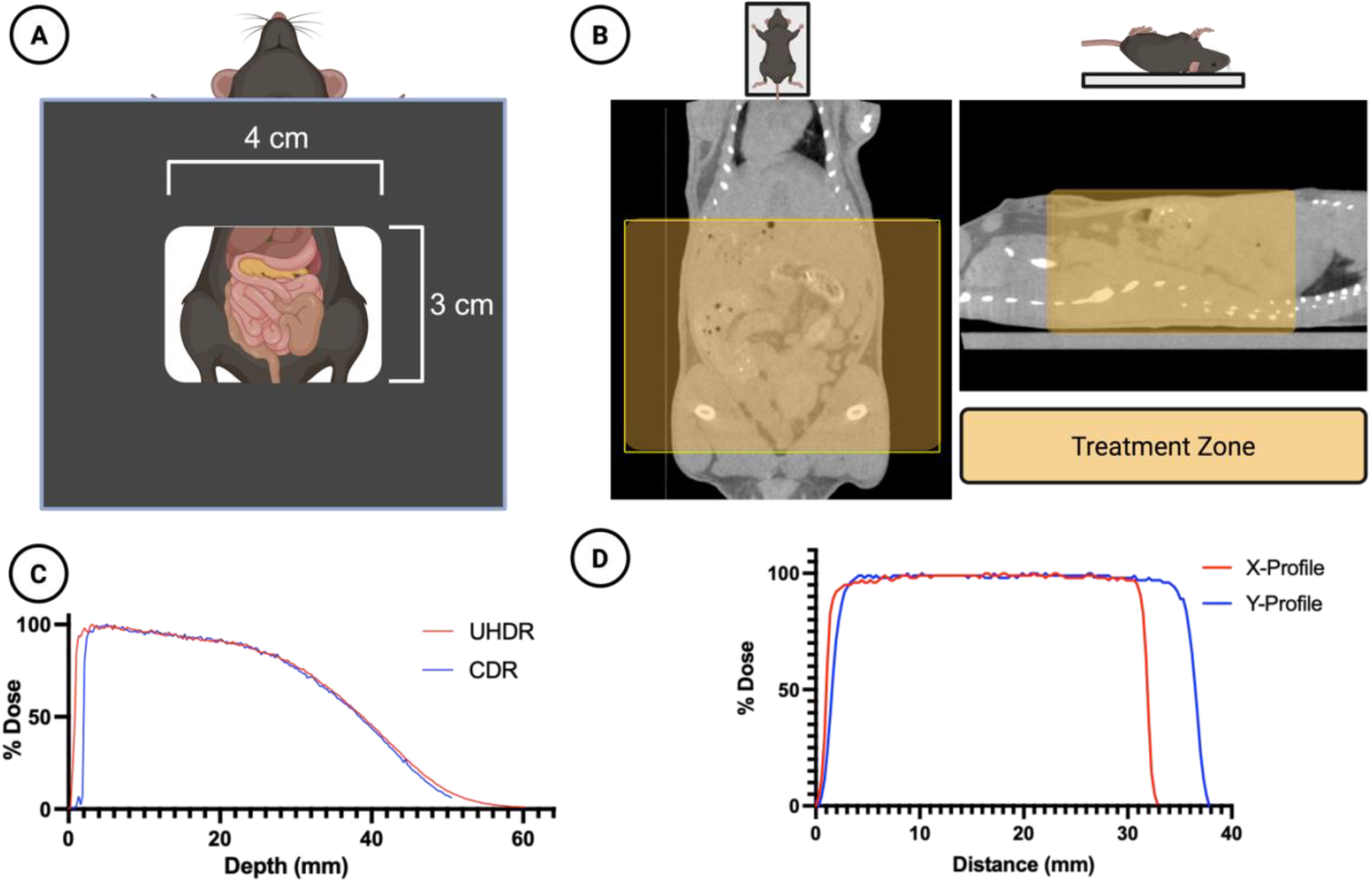
Treatment Design. (**A**) A 4×3 cm rectangular cutout was used to irradiate the mouse abdomens. (**B**) A micro-CT scan of the mice demonstrates the treatment zone (shaded yellow) superimposed in two planes. (**C**) The dose-depth characteristics of the cutout for both UHDR and CDR beams are shown in solid water for a 14Gy prescribed dose, obtained using the PTW FLASH diamond detector (**D**) The horizontal and vertical dose profiles of the irradiation field are shown, obtained using EBT-XD radiochromic film.

On the day of live animal irradiation, quality assurance was conducted to verify machine output in CDR and UHDR mode was within 3% of verification day. The calculated output was then inputted into the treatment planning program to determine the final treatment parameters. The final treatment parameters were delivered to the pre-calibrated uD detector at a depth of 1 mm in solid water and dose delivered was once again verified to be within 3% of the expected dose. Finally, individual dose delivery was monitored by placing the uD at the edge of the irradiation field.

### Radiation Damage Assay

Following irradiation mice were weighed daily and percent weight loss was calculated from the pre-irradiation baseline weight. Time to 25% weight loss was used as the primary endpoint for time to event (survival) analysis.

### Oxygen Measurements

Six (N=6) C57BL/6 mice received IV injections of PdG4 Oxyphor. The mice were subsequently anesthetized using either IP injection of Ketamine/Xylazine (ketamine 100 mg/kg, xylazine 10 mg/kg, n=3) or inhaled isoflurane anesthesia delivered in room air through a non-rebreather mask (3% induction for 3 minutes, then maintained on 1.5%, n=3). Upon verification of the surgical plane of anesthesia, a laparotomy was done to expose the abdominal cavity. Oxygen measurements were taken using the OxyLED (Oxygen Enterprises Inc, USA), with the optical fiber centered at the abdominal cavity.

### Statistical Analysis

Time to event data were analyzed using log-rank tests. Mice were censored in analyses if death occurred prior to 25% weight loss or if the weight loss was not achieved by 14 days post-irradiation. Oxygen measurements were analyzed using two sample t-tests. All analyses were conducted in Prism (GraphPad Software LLC., USA) with a significance level being α < 0.05.

## Results

### Weight Loss

Mice were anesthetized with ketamine/xylazine (K/X) or isoflurane, and irradiated with 14 Gy or 16 Gy of either UHDR or CDR. We used time to 25% weight loss as our primary survival endpoint. Following 14 Gy abdominal irradiation, 14 animals (8 in the UHDR K/X arm, 4 in the CDR K/X arm, and 2 in the CDR ISO arm) never reached the weight-loss threshold before the 14-day study cut-off and were therefore censored. Kaplan–Meier analysis showed that UHDR K/X mice maintained body weight significantly longer than UHDR ISO mice. Median time to endpoint was 6 days for UHDR-ISO and CDR-ISO, whereas it was 7.5 days for CDR-K/X and was not reached for UHDR-K/X, with 80 % of those animals still on study at day 14. This is demonstrated in Figure 1.

Following 16 Gy irradiation, 4 animals in the UHDR K/X arm did not reach the weight loss threshold before the study cut off and were censored. Kaplan–Meier analysis showed that UHDR K/X mice maintained body weight significantly longer than CDR K/X mice, whereas the ISO mice did not differ in weight. Median time to endpoint was 6 days for UHDR-ISO, CDR-ISO, and CDR K/X. UHDR K/X mice had a median time to endpoint of 7 days, but 40% remained on study at day 15. This is demonstrated in Figure 2.

**Figure 2:**
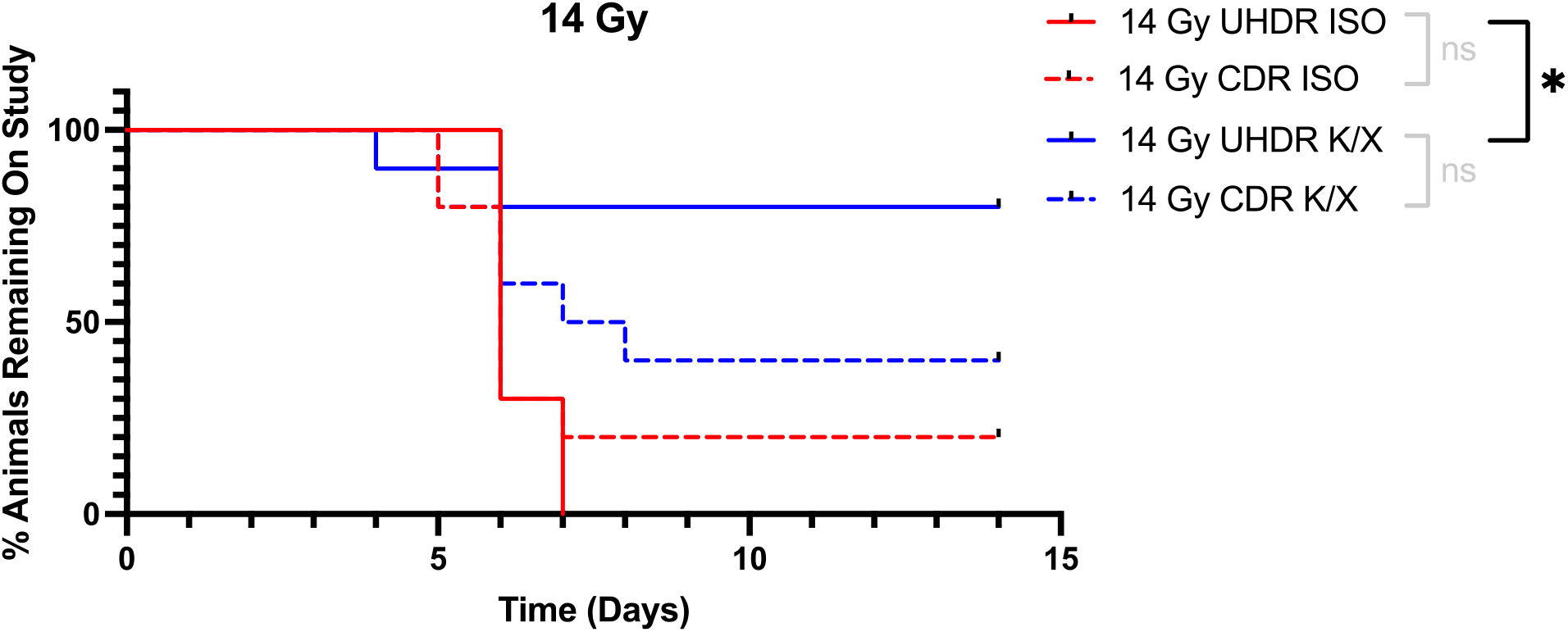
Kaplan-Meier curves for mice irradiated with 14 Gy. Mice were anesthetized with either isoflurane (ISO) or ketamine/xylazine (K/X) and irradiated with 14 Gy of conventional (CDR) or ultra-high dose rate (UHDR) irradiation. Mice were removed from study when they reached 25% weight loss from baseline. NS = not significant, * = p <0.05

**Figure 3:**
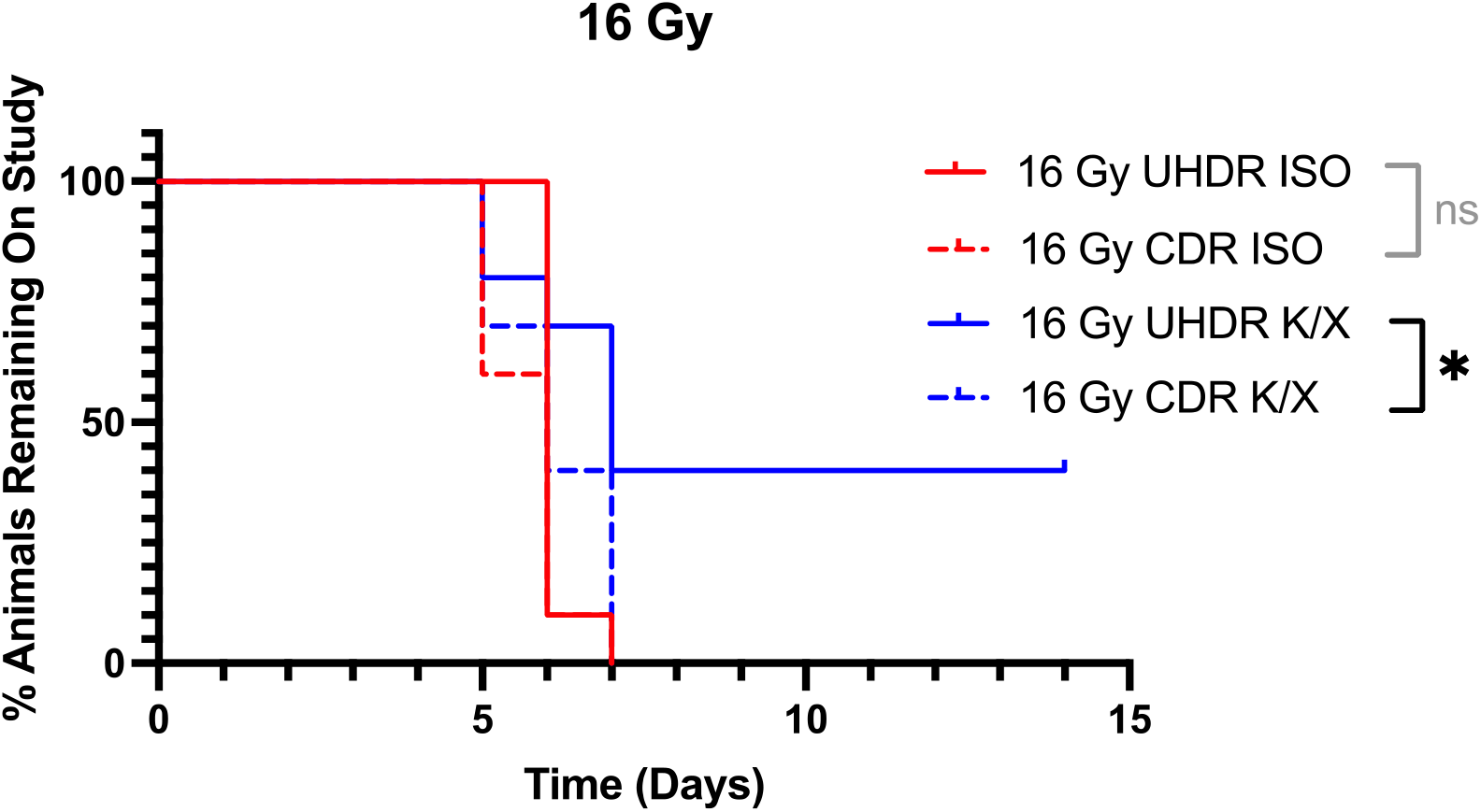
Kaplan-Meier curves for mice irradiated with 16 Gy. Mice were anesthetized with either isoflurane (ISO) or ketamine/xylazine (KX) and irradiated with 16 Gy of conventional (CDR) or ultra-high dose rate (UHDR) irradiation. Mice were removed from study when they reached 25% weight loss from baseline. NS = not significant, * = p <0.05

**Figure 4:**
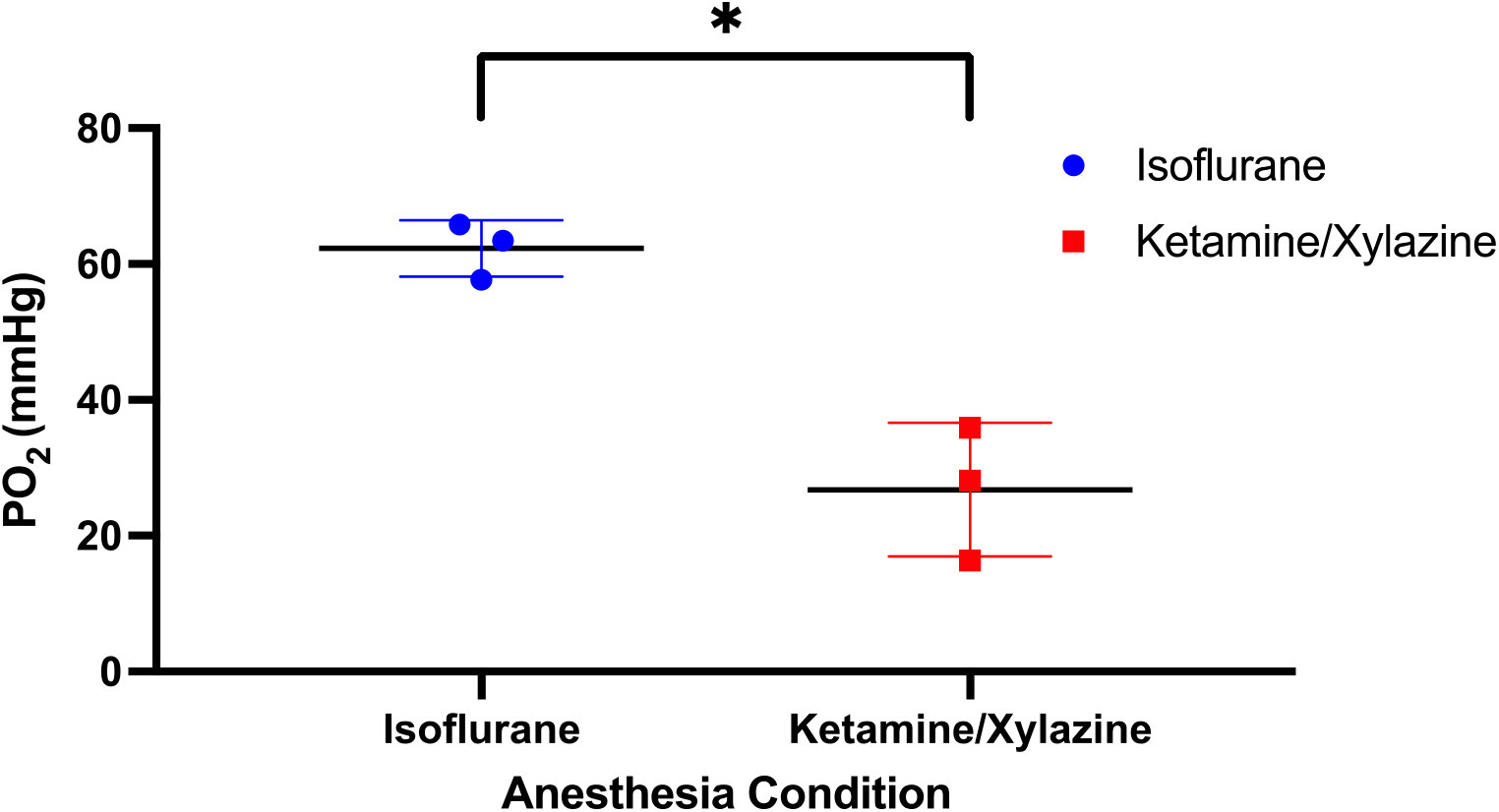
Abdomen oxygen measurements. Mice were anesthetized with either isoflurane or ketamine/xylazine, injected with PdG4 Oxyphor, and a laparotomy was done to expose the abdominal cavity. Oxygen measurements were obtained using the OxyLED system. * = p <0.05

### Oxygen Measurements

Bowel oxygen measurements through a laparotomy following systemic administration of PdG4 Oxyphor demonstrated significantly higher oxygen levels in mice anesthetized with isoflurane (mean=62 mmHg, SD=4) compared to mice anesthetized with ketamine/xylazine (mean=27 mmHg, SD=10).

## Discussion

Using a 9 MeV electron beam, two doses were delivered, at 14 or 16 Gy, with broad field irradiation to the abdomens of C57BL/6 mice under two commonly interchangeable anesthetics: (i) intraperitoneal ketamine/xylazine (K/X, 100/10 mg kg^−1^), and (ii) inhaled isoflurane (ISO, 3 % induction, 1.5 % maintenance) in room air. Supplemental oxygen was deliberately avoided to isolate anesthetic-specific physiology. Employing time-to-25% weight-loss as an objective endpoint, we asked whether the magnitude of FLASH depended on the anesthetic type.

At the 14 Gy dose, a robust radioprotective effect was seen in UHDR mice anesthetized with K/X compared to ISO. Though a statistically significant sparing effect was not seen between K/X UHDR and CDR mice, it is apparent that this may have been caused by limited statistical power from the few events in the UHDR K/X arm. Indeed, at termination of study (14 days), 80% of UHDR K/X arm had not reached 25% weight loss, compared to 40% in the CDR K/X arm. Importantly though, there is no apparent FLASH effect colon sparing in mice anesthetized with ISO, indicating that with ISO anesthesia this would not be seen at this dose level.

At the increased 16 Gy dose, K/X administration showed notable radioprotection to the UHDR irradiated mice, as compared to the CDR mice. A significant sparing effect was seen when comparing UHDR to CDR irradiated mice under the K/X condition as well. At the termination of study, 40% of the UHDR K/X mice had not reached the 25% weight loss endpoint, in comparison to CDR mice who all reached the endpoint by 7 days. These results support the conclusion that ISO anesthesia negates the FLASH sparing sparing effect at this dose level.

To investigate a potential mechanism for loss of sparing under ISO anesthesia, intraperitoneal oxygen measurements were performed using an injected optical oxygen reporter and fiber measurement system in six mice. A significant, nearly 2.5-fold difference in oxygen tension was seen between ISO (average ∼62 ± 4 mmHg) and K/X (∼26 ± 10 mmHg) conditions. Considering this in the context of recently published work investigating UHDR sparing under varying oxygen conditions in mouse skin, this data supports the interpretation that an oxygen-based mechanism affects the observed differences in radiosensitivity between ISO and K/X anesthesia. We see, again, that UHDR sparing is particularly sensitive to tissue oxygenation, beyond what would be expected from our conventional understanding of oxygen radio sensitization and the oxygen enhancement ratio.

There is now a wide body of literature demonstrating that tissue oxygenation is a key determinant of the magnitude of UHDR sparing across organs. This data suggests that an oxygenation threshold or spectrum must exist, above which FLASH sparing is not significant. It is widely known that oxygen enhanced damage through peroxyl formation and DNA damage fixation, and so higher oxygen contributes to higher damage, but this is only seen in the range of local pO2 <10 mmHg. Thus, it seems plausible that with UHDR irradiation that some local depletion occurs or local change in reactive oxygen species occurs, which only significantly affects the normal tissues with lower initial pO_2_. questions remain about what this threshold is, whether it is organ-specific, and if it is modulated by total dose, dose-rate, or fractionation. Once these questions are answered, we can then tackle what we consider to be the key oxygenation question in translational UHDR: Are physiological oxygen values in awake, normally ventilating humans low enough to see a clinically meaningful UHDR sparing effect?

Our study is not without limitations. First, although we see a potent sparing in in mice who were anesthetized with K/X and received 14 Gy of UHDR radiation compared to CDR, we fail to detect a statistically significant difference using Kaplan-Meier (KM) analysis. This isattributed to the limited number of mice meeting the endpoint (2) in the UHDR arm, which affords very little statistical power to the log-rank test. Still, considering the two dose levels 14 Gy and 16 Gy together, UHDR sparing is most apparent under the K/X condition across doses. The second limitation is the inherent uncertainty in the abdominal compartment that is probed in our oxygen measurements. In Total Abdominal Irradiation (TAI), numerous organs are irradiated with vastly different radiosensitivities. Though weight loss following TAI is typically attributed to small bowel damage, the more nuanced reality is that damage to many of the intrabdominal structures, aside from small bowel, can lead to weight loss. In post-mortem gross necropsy, significant hydronephrosis, liver damage, and gastric enlargement were noted in addition to enteritis. The wide-field oxygen measurements performed centered on the bowel to maximize signal from this organ of interest. This means that these oxygen values are an average of numerous abdominal structures within the irradiation field, rather than any one organ, but were weighted toward the bowel. Still, the results are clear: intrabdominal oxygen levels under ISO anesthesia are significantly higher than under K/X anesthesia. As oxygen measurement technologies advance, more organ-specific measurements would add value to TAI studies.

In summary, by explicitly integrating anesthetic physiology, dosimetry verified to within 3 %, and direct oxygen measurements, the present work identified one of the persistent sources of variability in pre-clinical UHDR research, which is anesthesia method. There needs to be practical guidance for experimental design as the field advances toward FLASH clinical trials. While direct measurement of tissue oxygen is challenging, future investigations should consider tighter control and reporting of experimental factors that can alter tissue oxygenation.

